# Parasite mediated competition facilitates invasion

**DOI:** 10.1101/167254

**Authors:** Senay Yitbarek, Ivette Perfecto, John H. Vandermeer

## Abstract

Parasites play an important role in invasion success with important consequences for biodiversity and community structure. While much research has focused on direct effects of parasites on biological invasions, parasites can also indirectly influence interactions within the invaded community across trophic levels. For instance, parasites can mediate competitive interactions between native and exotic species through trait-mediated indirect effects. We consider the interactions between the parasitoid fly Pseudacteon sp. (Diptera: Phoridae), and its native host ant Linipethema iniquum, and the exotic ant Wasmannia auropunctata in the introduced range of Puerto Rico. We examined the effects of phorid flies on the competitive outcome between the arboreal ants W. auropunctata and native ant L. iniquum. Furthermore, we investigate the searching efficiency of phorid flies in detecting L. iniquum nests. To study the indirect effects on ant competition, we monitored ant recruitment to baits over a 60-min time interval in the presence and absence of phorid fly parasitoids. We then performed field experiments and measured phorid arrival time to arboreal nests of L. iniquum located in both a) W. auropunctata patches and in b) isolated patches dominated by L. iniquum nests. We found that the presence of phorid fly significantly reduced recruitment of L. iniquum workers to baits through induced behavioral changes thereby increasing the ability of W. auropunctata to acquire resources. In addition, we found that phorid arrival time in isolated patches of L. iniquum patches was faster as compared to L. iniquum nests located within W. auropunctata patches. Our results show that phorid fly parasitoids indirectly may influence competitive interactions by attacking the host-ant L. iniquum and consequently providing an advantage to local spread of W. auropuntata populations in Puerto Rico. However, the spatial dynamics of arboreal ants shows that L. iniquum seeks protection from phorid fly parasotoids by moving their nests to W. auropunctata dominated patches.

## 1 Introduction

Biological invasions pose a major challenge to ecosystems with consequences for community structure and biodiversity (Dunn et al. 2012). The global spread of introduced species is altering native communities and is rapidly accelerating due to human activities (Mooney and Cleland 2001). In recent years, there has been growing interest in the role of parasites in invasion success (Tompkins et al. 2010). Much of the research involving the role of parasites in invasions has focused on direct interactions, including competition and predation studies, and on their subsequent effects on community dynamics (Wilson et al. 1998, Corbin and D’ Antonio 2004). However, indirect effects are believed to play a key role in the structuring of ecological communities (Holt 1977, Miller 1994, Bonsall and Hassell 1998, Peacor and Werner 2001). The extent to which indirect effects influence the invasion process and their consequences on native communities remains unexplored (White et al. 2006). Parasites can indirectly structure ecological communities within the same trophic level through parasite mediated competition (Bowers and Turner 1997). During biological invasions, the absence of parasites can indirectly enhance the competitive ability of an exotic species (Torchin et al. 2003) by reallocating resources against defense to competitive traits (Blossey and Notzold 1995). Parasites can affect the outcome of competitive interactions between exotic and native species through both density-mediated indirect effects (DMII) and trait-mediated indirect effects (TMII). For instance, competitive interactions between the native ant Solenopsis geminata and invasive ants Solenopsis. invicta were altered by the phorid parasitoid Pseudacteon browni (Morrison 1999). In the presence of phorid flies, S. geminata had to defend themselves against P. browni resulting in 50 % decline in resource retrieval thereby giving the invasive ant S. invicta a competitive advantage. As such, Pseudacteon phorid fly parasitoids (Diptera: Phoridae) have been used as a biological control agent against S. invicta populations (Mehdiabadi et al. 2004). Interspecific competitive trade-offs are believed to play an important role in the structuring of ant communities (Lebrun and Feener 2007). The discovery-dominance trade-off describes the ability of a species to discover a food resource versus the ability of a species to dominate a food resource (Fellers 1987, Savolainen et al. 1988). The discovery-dominance trade-off allows for species coexistence and has been documented in studies involving only a few ant species (Lynch et al. 1980, Perfecto 1994, Morrison 1996, Feener et al. 2008). However, exotic ants are believed to break down this trade-off in their introduced range by excelling at both traits (Holway 1999). Another trade-off involves the ability of species to defend itself against natural enemies in contrast to maximizing their competitive abilities (Lebrun and Feener 2007). For instance, specialized phorid fly parasitoids have been found to attack host ants and limit their foraging activities resulting in the frequent loss of resources to their competitors (Feener and Brown 1992) (Orr et al. 1995, Morrison et al. 2000, Philpott 2005). Together these trade-offs interact with one another to influence community structure in such a way that a species ability to maximize it’s competitive potential is balanced by its vulnerability to parasitism (Adler 1999). An optimal fitness strategy is to minimize competitive ability to the level of the entire ant assemblage in order to avoid parasitism (Adler et al. 2007). While the presence of phorids has been found to reduces foraging rates to baits, phorids may not always determine the outcome of competitive interactions between native and exotic ant species (Morrison 1999). In addition, phorids were not influential in competitive outcomes between Solenopsis invicta and Solenopsis geminata in a laboratory setting (Morrison 2000). Furthermore, the spatial distribution of hosts can affect the searching efficiency and attack rates by phorids (Philpott et al. 2009). Empirical studies are needed to address the relative importance of phorid parasitoids in influencing invasion dynamics and community structure. Our study focuses on the invasion dynamics of W. auropunctata and competitive interactions with the arboreal ant Linipethema iniquum in Puerto Rico. W. auropunctata is native to Central and South America and has in recent decades expanded to island groups in the Carribean and Pacific Oceans (Foucaud et al. 2010). It has also spread to parts of Western Africa, including Gabon and Cameroon, and most recently to the Middle East (Walker 2006) (Ndoutoume-Ndong and Mikissa 2007, Vonshak et al. 2009, Mikkisa et al. 2013, Wetterer 2013). Within its native range, W. auropunctata is regarded as a common, but sub-dominant species as it faces intense competition by dominant ant species (unpublished data). W. auropunctata is widely distributed in Puerto Rico and can reach high populations densities on coffee farms (Wetterer 2013). At our site, phorid parasitoids were found to be attacking L. iniquum workers (Pseudacteon sp.). This study addresses whether trait-mediated indirect effects by phorids mediate competition and facilitate the invasion of W. auropunctata in Puerto Rico (fig 4.1). We examined the effects of phorid parasitoids on resource competition between host L. iniquum and non-host W. auropunctata and the effects of the spatial distribution of ants on phorid searching efficiency.

**Figure 1.**
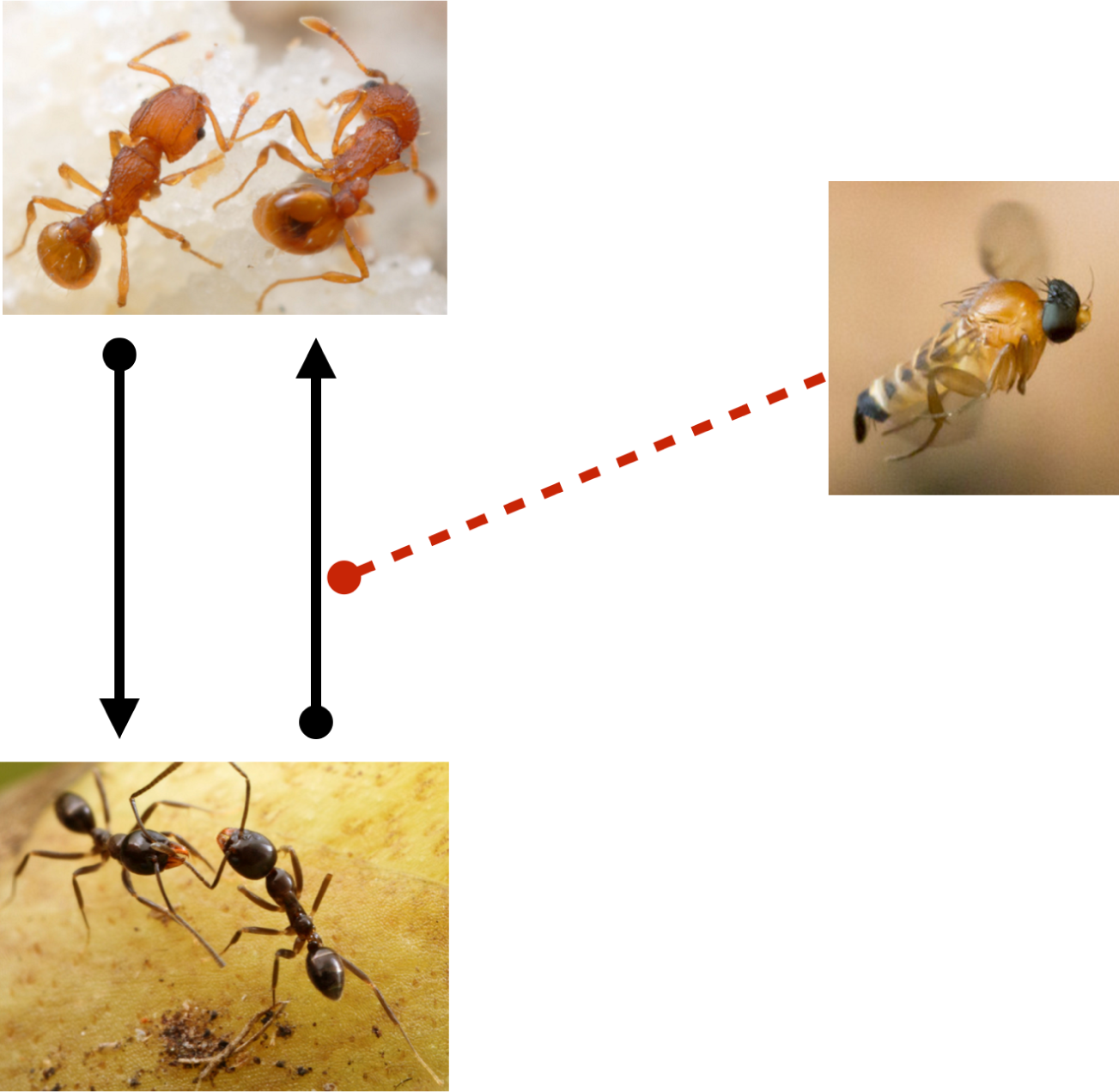
Diagram indicating direct effects (black solid lines) and trait-mediated indirect effects (red dashed lines). Competitive interactions between *Wasmannia auropunctata* (top picture) and *Linipethema iniquum* (bottom picture) are mediated by parasitoid phorid flies. Photo credits: Alexander Wild

## 2 Materials and Methods

### 2.1 Study sites and species

The study was conducted in the months of February, June, and July, in 2015 and 2016, on a coffee farm in Puerto Rico. The field site consisted of a 5-hectare plot within a high-shade organic coffee farm located in the central mountainous region in the municipality of Orocovos (18.175850, −66.4155700). The plot consisted primarily of coffee and banana trees where W. auropunctata and L. iniquum were found to be nesting. W. auropunctata invasion into the Carribean region is the result of multiple introductions due to human-induced dispersal and likely originates from it’s native habitat along the northern coast of South America (Foucaud et al. 2010). The Caribbean region represents an important zone from where secondary dispersal that led to the global expansion of W. auropunctata. The native range of L. inipethema extends from South to Central America into the eastern parts of the Caribbean (Wild 2008). L. iniquum is an arboreal ant species and has been found nesting in hollow twigs and leaf petioles of plantain in Puerto Rico (Wheeler 1908). We surveyed the 5-ha plot to determine the abundance and spatial distribution of W. auropunctata and L. iniquum populations (fig 4.2). Tuna baits were placed along transects of coffee and banana trees spaced out every 2 meters. Baits were subsequently checked every 30 minutes to determine the presence or absence of W. auropunctata and L. iniquum. In order to examine competitive interactions a small plot (30 m × 14 m) was established along the edges of W. auropunctata and L. iniquum territorial boundaries. A total of 324 baits were placed on trees in the large plot (within the high-shade organic farm). Surveys were completed in June and July of 2015.

**Figure 2.**
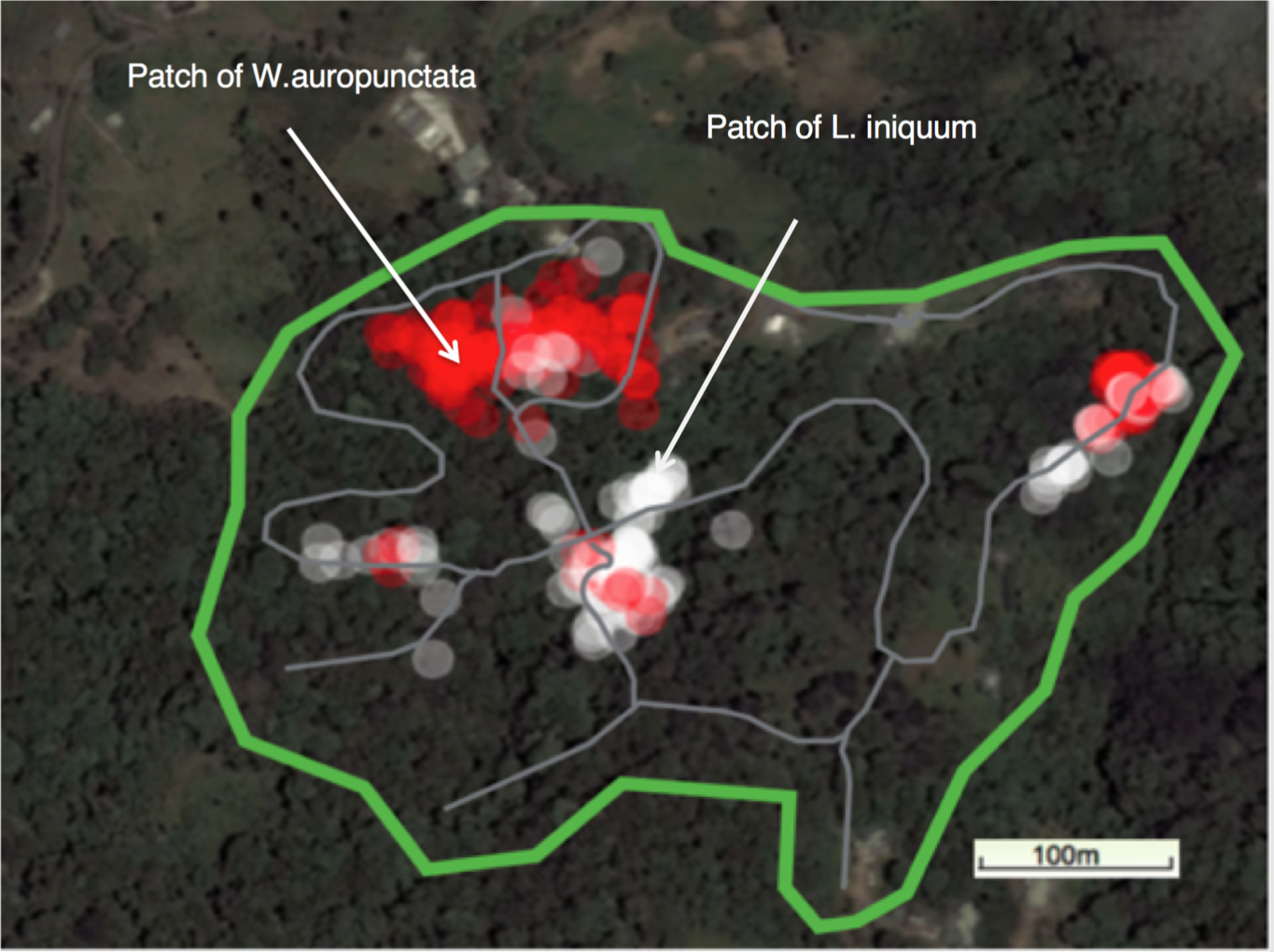
Spatial distribution of *Wasmannia auropunctata* (red) and *Linepithema iniquum* (white/gray) arboreal nests over a 7-ha plot in Puerto Rico. *W. auropunctata* nests dominate the largest patch (1-ha). Individual trees with *W. auropunctata* are scattered within smaller clusters of *L. iniquum* nests, while in the *W. auropunctata* patch several trees are occupied by *L. iniquum*.

### 2.2 Effects of parasitoids on ant competition

To test the trait-mediated indirect interactions by phorid flies on ant interactions we experimentally 1). Quantified the number of ant workers competing over time in the presence and absence of phorid flies. 2) Transplanted nests in trees dominated by W. auropunctata and L. iniquum. To assess competitive ant interactions, colonies of W. auropunctata and L. iniquum were randomly selected from coffee trees in the plot. To attract and collect phorid flies in the field, workers of L. iniquum were squashed by hand at each tree stand. Phorid flies were aspirated and placed in plastic containers in tents. Due to their fragility, phorid flies were kept in containers for less than 2 hours. Controls consisted of collected W. auropunctata and L. iniquum nests placed in containers connected by an artificial platform with droplets of honey resources placed at the center. Treatments consisted of W. auropunctata and L. iniquum colonies, in addition to phorid flies that were introduced at the beginning the experiment. Once the ants began foraging and replenished with honey, we recorded the number of ants at the bait ever minute for a maximum of 60 minutes or until all baits were occupied by 10 or more ants. Experiments were replicated 4 times for each interaction.

### 2.3 Parasitoid searching efficiency

We measured arrival times of phorid flies to trees occupied by L. iniquum. We identified four major ant clusters within the 5-ha plot. The overwhelming majority of trees were occupied by W. auropunctata, while the remaining trees were dominated by L. iniquum and Brachymyrmex species (fig 4.2). Within the largest W. auropunctata cluster, several trees were occupied by L. iniquum. Within the three smaller clusters, W. auropunctata was detected on several trees. We randomly selected four trees occupied by L. iniquum in the largest cluster and four trees occupied in the smaller clusters. At each tree, we disturbed L. iniquum nests by squashing their workers by hand and recorded the arrival time of phorid flies.

### 2.4 Data analysis

To compare recruitment of W. auropunctata and L. iniquum to baits with and without phorids, we used fisher’s exact test to compare the number of ant workers recruited over time to baits, testing the null hypothesis that the number of foragers is the same between treatments. Each treatment consisted of W. auropunctata and L. iniquum colonies competing for tuna baits resources in the presence of phorids, while in the controls phorids were excluded. To investigate the effects of the spatial distribution of arboreal ants on phorids, we examined the arrival time of phorids to trees dominated by L. iniquum. We used a simple t-test to determine differences in phorid arrival time between sites that included L. iniquum trees within clusters dominated by W. auropunctata and sites with trees dominated by L. iniquum with W. auropunctata trees interspersed (fig 4.2).

### 3 Results

Phorids flies influenced the competitive interactions between W. auropunctata and L. iniquum. We observed that L. iniquum in the presence of phorids underwent a 3-fold reduction in less than 15 minutes, which caused significant declines of L. iniquum abundance (p = 0.01). W. auropunctata abundances significantly increased abundance levels by 2-4 times in the presence of phorids (p < 0.0001). Phorid flies limited the recruitment rate of L. iniquum (fig 4.3), likely restricting resource uptake, which in effect increased the abundance of W. auropunctata in the presence of phorids. The spatial distribution of L. iniquum nests in trees significantly affected the searching efficiency of phorids. The majority of trees where occupied by W. auropunctata nests (N=166), followed by L. iniquum nests (N=129), and unidentified ant species (N=29). We found a significant effect of phorid arrival in L. iniquum dominated patches as compared to W. auropunctata dominant patches. Phorids in nearby L. iniquum patches took anywhere between 3-5 minutes to arrive at trees, while phorids took much longer to detect L. iniquum trees within W. auropunctata patches (fig 4.4). Thus, the spatial distribution of L. iniquum limited the effectiveness of phorid control in areas where W. auropunctata dominates.

**Figure 3.**
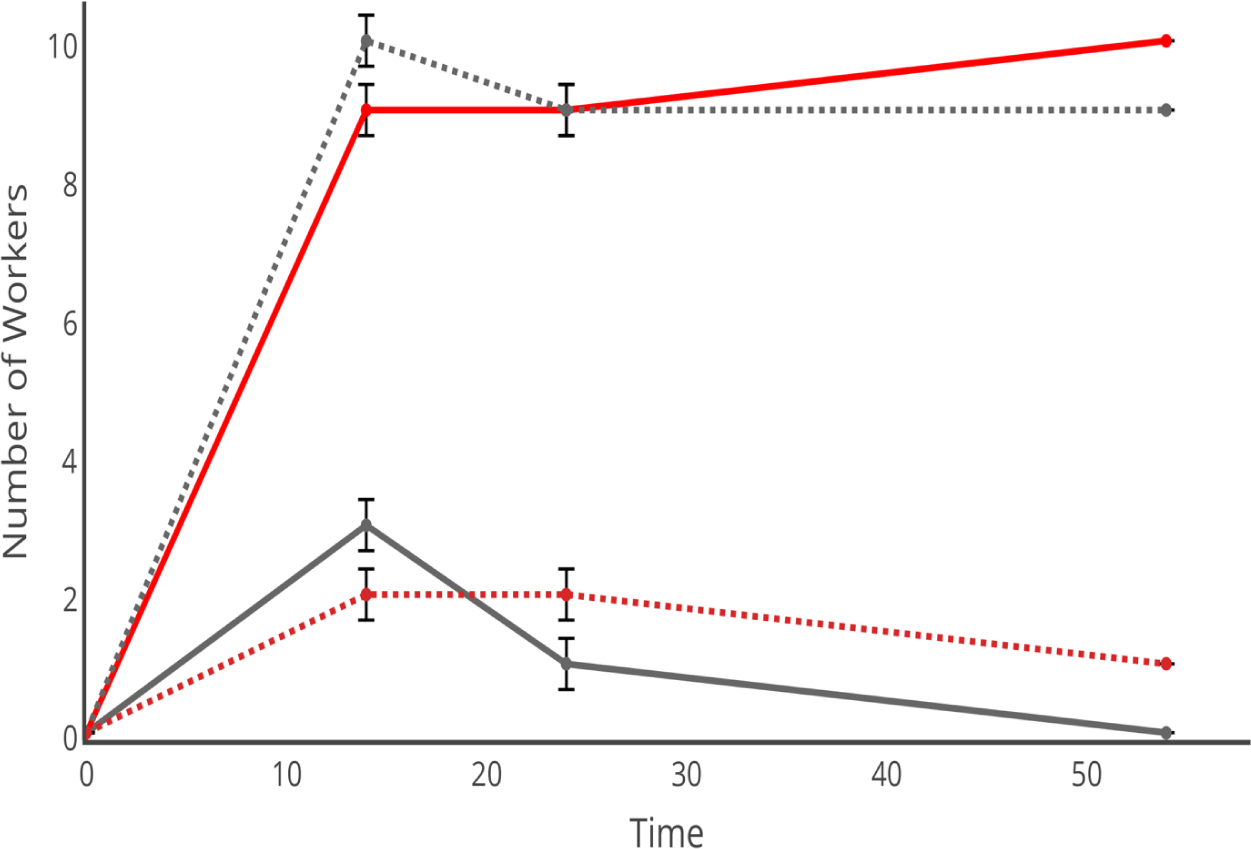
Arboreal ant foraging dynamics in the presence and absence of phorid flies. In the absence of phorid flies, the native ant *L. iniquum* (grey-dashed line) out-competes the invader *W. auropunctata* (red-dashed line). In the presence of phorids, L. iniquum (solid grey) workers rapidly decline while the number of W. auropunctata (solid red) remains relatively constant (P < 0.0001).

**Figure 4.**
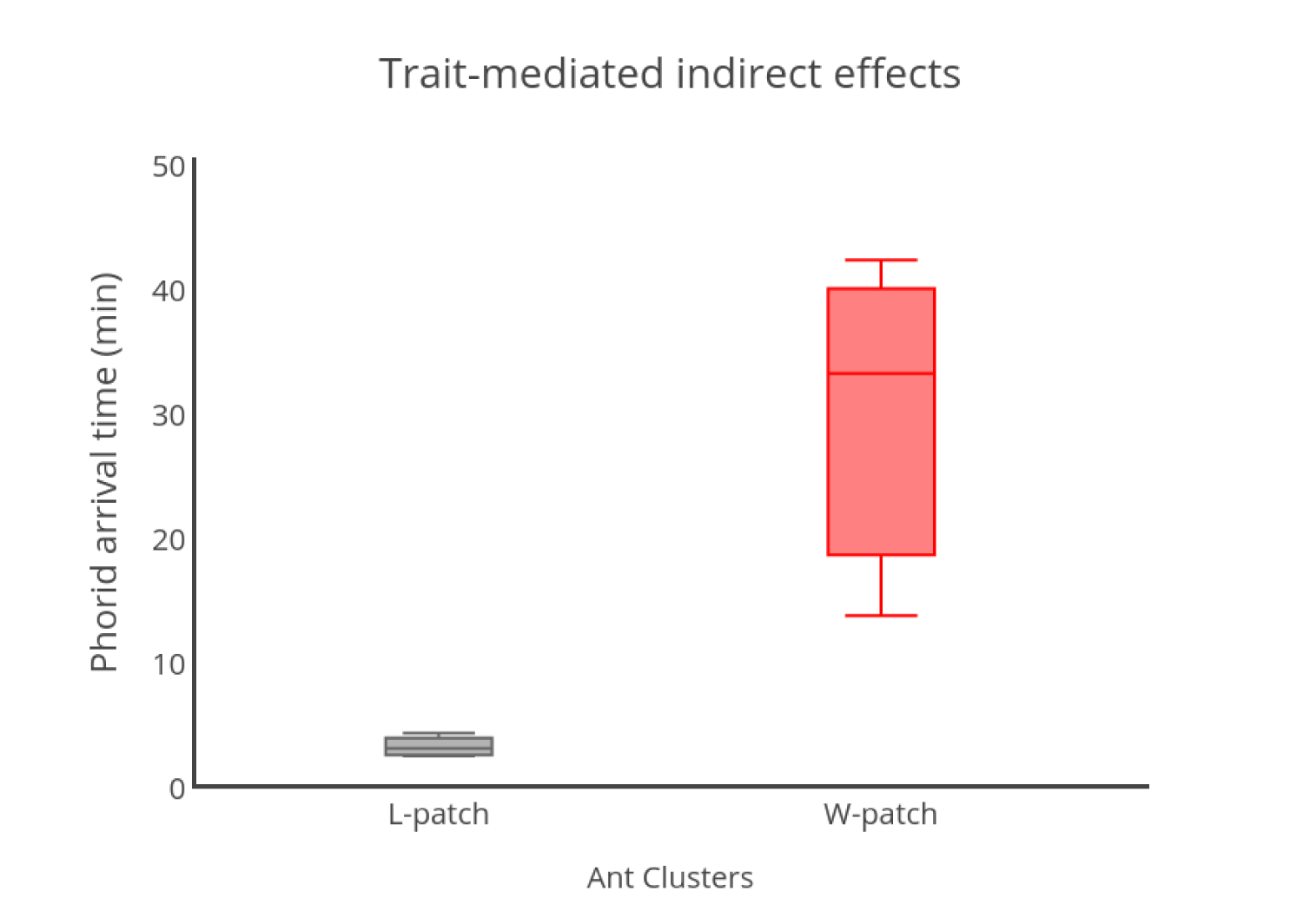
Phorid searching efficiency of *L. iniquum* hosts in the field. Phorid arrival time was faster in *L. iniquum* dominated patches (grey-box) as compared to *L. iniquum* found in W. auropunctata patches (red-box), (t=-3.14, df=4, p=0.03)

### 4 Discussion

Phorid fly presence significantly affected competitive interactions between *W. auropunctata* and *L. iniquum*. Phorid flies significantly affected the recruitment of *L. iniquum* workers to tuna baits, which significantly increased the overall abundance levels of *W. auropunctata* workers. In the absence of phorid flies, *L. iniquum* workers had a higher recruitment rate to baits and gained a competitive advantage over *W. auropunctata*. In the presence of phorids, *W. auropunctata* workers were able to overtake baits through a combination of direct interactions (i.e. competition) and indirect attacks by phorids on *L. iniquum*. This double-whammy led to the rapid decline of *L. iniquum* workers over time. The effect of phorids on ant community structure depends on the spatial scales of interaction. Our results indicate that phorid searching efficiency depends on the spatial scale of both the phorid fly and distribution of *L. iniquum* nests. It took much longer for phorids to locate L. iniquum nests within the patch of W. auropunctata as compared to areas where L. iniquum was more prevalent. As distance increased from densely populated areas of L. iniquum, phorid attack rates diminished because of the time it took to locate nests. Although we don’t claim to have information on the population density of phorids at local spatial scales, it appears that phorid flies increase their local population densities nearby areas dominated L. iniquum. Phorid flies are known to use chemical cues to locate their hosts (Brown and Feener 1991, Morrison 1999, LeBrun and Feener 2002, Hsieh and Perfecto 2012). During competitive encounters, ants release alarm pheromones resulting in the rapid response by phorid flies. One study found that phorid attack rates increased when Linipethema was in the presence of dominant ant competitors (Orr et al. 2003). Thus, a behavioral response by Linipethema during competitive encounters elicits a positive feedback by increasing phorid attack rates which increases the number of foragers of its opponents (Hsieh and Perfecto 2012, Philpott 2005). At larger spatial scales, we found that phorid attack rates diminished at greater distances from L. iniquum clusters. Interestingly, L. iniquum nests found within W. auropunctata clusters were temporarily relieved from phorid attacks. There was a five-fold reduction in phorid attack rates in isolated L. iniquum trees within W. auropunctata clusters which suggests that L. iniquum species disperse to patches dominated W. auropunctata in order to receive protection against phorid parasitoids. The effects of phorids in mediating ant competition may vary temporally. Our results demonstrate that L. iniquum abundances were reduced in the presence of phorids during the experiment. However, ants can adapt the timing of their foraging activities as a way of avoiding phorid attacks (Philpott et al. 2004). For example, the ground forager ant Atta cephalotes primarily forages during the night time in order to avoid phorids during the day time that tend to build up their populations outside of A. cephalotes nest colony entrances (Orr 1992). Several Linipethema species construct mobile nests that are often disguished from phorids making it more difficult to find (Markin 1970). In our system, L. iniquum species have many different shallow nest entrances throughout coffee trees making it even more difficult for phorids to figure out where to attack. Therefore, species differences in nesting ecology and habitat complexity may play an important role in phorid attacking abilities (Orr et al. 2003) (Philpott 2005). Another important factor in assessing TMII on competition is resource size (Mehdiabadi et al. 2004, Philpott et al. 2004). In our study, both L. iniquum and W. auropunctata were provided with large resources. However, this may not have given L. iniquum enough time to defend itself from phorids as they recruit large number of workers to uptake resources. In our experiment, L. iniquum was rapid at resource discovery, as compared to W. auropunctata, but spent longer time recruiting workers to carry baits. Although not enough is known about species response to variation in resources, small resources distributed over larger spatial scales may enable L. iniquum to avoid persistent phorid attacks. (LeBrun 2004) showed that phorids take a much longer time locating ants consuming smaller resources. As fewer ants are recruited to smaller resources, less pheromones are being released that phorids can exploit to detect ant foraging activities (Folgarait and Gilbert 1999) (Feener and Brown 1992). These findings have important implications for the invasion dynamics of W. auropunctata in a coffee agricultural ecosystem. In the absence of phorids, L. iniquum reaches higher abundance levels resulting in increased access to resources. This pattern changes however when phorids come into the scene and L. iniquum abundance diminish rapidly enabling W. auropunctata to increase its abundance levels. At the population level, L. iniquum builds up locally dense clusters but undergoes increased attacks by phorids, which provides an opportunity for W. auropunctata to invade. To counter this two-fold attack, L. iniquum disperses into patches dominated by W. auropunctata, providing temporary protection from phorid attacks. These factors taking together show clear effects of phorids on ant competition. Studies of ant competition have shown the existence of competitive trade-offs among species can structure ecological communities and promote coexistence (Fellers 1987, Savolainen et al. 1988, Perfecto and Vandermeer 2013). The trade-off implies that some species have strong resource discovery abilities as opposed to other species that have strong resource domination abilities. TMII mediated competitive interactions by phorids can break the discovery-dominance trade-off and enable invasive species to spread (Hsieh and Perfecto 2012). In the absence of phorids, L. iniquum is a faster resource discoverer and gains a competitive advantage over W. auropunctata species, which are known to be poor resource discoverers (Bertelsmeier et al. 2015). However, phorids can tip the balance and limit the rapid acquisition of resources by L. inquum thereby providing W. auropunctata species with ample time to dominate resources and spread. A similar pattern was found in a pine-oak woodland ecosystem involving Pheidole host species that in the absence of phorids increased their discovery abilities, thereby breaking the discovery-dominance trade-off. The presence of phorids reduced the competitive ability of the host species to the level of the ant assemblage reinforcing the discovery-dominance trade-off (Lebrun and Feener 2007). Phorid incuced TMII attacks on L. iniquum enable to W. auropunctata to spread in Puerto Rico. A key question that remains is what potential factors limit the expansion of W. auropunctata in its native range. To our knowledge no phorid parasitoids have been recorded in the native range of W. auropunctata. However, previous research in the native range of Mexico suggests that dominant ant species can limit the expansion of W. auropunctata in a Mexican agricultural ecosystem, providing further support for the biotic resistance hypothesis (Yitbarek et al. in press). In the case of another invasive ant species S. invicta, phorid induced TMII and interspecific competition influenced the ecological success of the invasive ant S. invicta (Feener et al. 2008). However, this pattern was found to vary geographically depending on the presence of phorid parasitoids in the region. In the case of Puerto Rico, we find that a combination of release from competitors in the native range and TMII contributes to the expansion of W. auropunctata. Our study on the role of phorid parasitoids mediating competitive ant interactions has important consequences for biodiversity maintenance. Competitive dynamics between W. auropunctata and L. iniquum for resources suggest that two different trade-offs operate. The ability of L. iniquum to arrive at resources quickly and the ability for W. auropunctata to hold on to acquired resources shows that a discovery-dominance maintains species coexistence in the absence of phorids. The discovery-dominance vanishes in the presence of phorids and in place a new trade-off emerges between the ability to compete for resource versus the ability to defend against parasitoids (Lebrun and Feener 2007). This so-called competition-defense trade-off reduced L. iniquum workers at resources while W. auropunctata was able to increase its workers to baits. The interplay between these trade-offs can amplify within complex ecological networks whereby the addition of a parasitoid reduces the dominance of a species to the level of the entire assemblage resulting in coexistence. From a practical standpoint, W. auropunctata has been found to be an effective biological control against the coffee berry borer pest in Mexico (Gonthier et al. 2013). Indirect effects by phorids enables W. auropunctata to expand in shaded areas with relatively high densities of the coffee berry borer. One potential mechanism for limiting the expansion of *W. auropunctata* on coffee farms is through the management of shade trees. Pruning of trees provides enough sunlight to attract ant competitors that can potentially limit the expansion of *W. auropunctata* while also providing ecosystem services. These findings open up a new set of inquiry on the effects of TMII on ant community structure across temporal and spatial scales. While phorids reduced the abundance level of *L. iniquum* resulting in a competitive advantage for *W. auropunctata*, we know very little about the long-term population level consequences of TMII on ant communities. In particular, it’s important to explore whether parasitoid phorids attacks on L. iniquum have implications for colony growth (Mehdiabadi et al. 2004). Although phorids reduce the number of workers of host species this may not necessarily affect the colony as a whole and therefore future investigations should evaluate the long-term dynamics between ants and phorids (Philpott 2005). Spatial considerations between ants and phorids need to take into account the foraging range of phorids. While phorids appear to build up their populations nearby L. iniquum clusters it’s not clear how far the range of phorids extends. Within W. auropunctata clusters several colonies of L. iniquum have been established in order to limit phorid attacks. This feedback between phorid range and ant competition leads to the formation of spatial clusters. In this regard, spatially explicit models can serve us useful guides to disentangle complex direct and indirect interactions between ants and phorids that give rise to spatial pattern formation and have important consequences for biodiversity (Li et al. 2016).

